# Investigating the effect of Ser256 phosphorylation on gating of aquaporin-2: Molecular Dynamics study

**DOI:** 10.1101/2021.01.18.427094

**Authors:** Pragya Priyadarshini, Balvinder Singh

**Author notes:** Correspondence: Dr. Balvinder Singh.

## Abstract

Regulation of water transport via aquaporins is crucial for osmoregulation and water homeostasis of an organism. This transport of water is regulated either by gating or trafficking wherein AQPs are transported from intracellular storage sites to plasma membrane. It has been proposed that water movement via AQP2 is regulated by post-translational modification. We aimed to explore the structural and functional changes occurring in AQP2 due to Ser256 phosphorylation. We have carried out molecular dynamics simulations to investigate molecular basis of effect of phosphorylation on water permeability of AQP2. MD simulations show that there are mild variations in the pore sizes of different monomers of the phosphorylated and unphosphorylated AQP2. Analysis of inter and intra-monomeric interactions such as hydrogen bond, electrostatic and hydrophobic interactions has been carried out. Structures of the phosphorylated AQP2 do not show any blocking of mouth of pore of the monomers during the course of MD simulations. Further, water permeability calculations do corroborate the above finding. This molecular dynamics study suggests that phosphorylation of C-terminal Ser-256 residue of AQP2 may not be directly responsible for gating mechanism.

## Introduction

Aquaporins (AQPs) have been well characterized as membrane channels that facilitate transport of water and small solute across the membrane (Borgnia et al., 1999; Soveral and Casini, 2016). Aquaporins are tetrameric protein, and a monomer is made up of six transmembrane α-helices and two short helices, with the amino and the carboxyl terminus located on the cytoplasmic surface of the membrane (Kruse et al., 2006). Each monomer individually can facilitate water flow through the channel. However, studies have shown that central pore of the tetramer in some AQPs is responsible for conduction of ions such as K+, Cs+, Na+,and tetramethylammonium (Saparov et al., 2001; Yool and Weinstein, 2002). All aquaporins have conserved NPA (Asn-Pro-Ala) motif near the center and aromatic/Arginine (ar/R) selectivity filter towards extracellular side in common and these two form constriction regions within the channel. AQPs participate in many physiological and pathophysiological processes that include renal water absorption, brain water homeostasis, fat metabolism, liver gluconeogenesis, tumor angiogenesis and reproduction (Soveral and Casini, 2016). Regulation of aquaporin’s function is crucial for osmoregulation and water homeostasis (Madeira et al., 2016). Water-selective AQPs are post-translationally regulated either by gating or trafficking where AQPs are transported from intracellular storage sites to the plasma membrane. Various factors like phosphorylation, intracellular Ca^2+^concentration, pH, pressure, solute gradient, temperature have been reported to affect this gating mechanism of aquaporin channel (Tornroth-Horsefield et al., 2010). AQPs can be subjected to regulation via different mechanisms, among which pH regulation has been disclosed for plant and for a few mammalian AQPs, such as AQP0, AQP3, and AQP6 (Nemeth-Cahalan and Hall, 2000; Yasui et al., 1999). Phosphorylation induced gating is well-studied phenomenon in SoPIP2;1, a spinach aquaporin. Different Molecular Dynamics (MD) studies of SoPIP2;1 show that phosphorylation at residues Ser115 and Ser188 leads into conformational changes in loop D affecting channel opening (Nyblom et al., 2009; Tornroth-Horsefield et al., 2006). The effect of phosphorylation has also been studied in rat AQP4 showing the insignificant effect of phosphorylation of Ser180 on water permeability although the interaction of phosphoserine-180 with C-terminal region causes movement of this C-terminal towards cytoplasmic mouth but it does not act as a lid for channel mouth. The osmotic permeabilities have been found to be comparable in case of phosphorylated and wild type rAQP4. Thus MD studies, do not support any exclusive gating effect of phosphorylation (Sachdeva and Singh, 2014). Nemeth-Cahalan and Hall have measured the rate of swelling of *Xenopus* oocytes injected with AQP0 cRNA in hypotonic solution under the effect of external pH and Ca^2+^. They found that reducing pH and Ca^2+^ in presence of extracellular His40 exclusive for AQP0, results into increase in water permeability (Nemeth-Cahalan and Hall, 2000). Nyblom et al. illustrates the molecular detail which adds into further details of channel gating in SOPIP2 (Nyblom et al., 2009). They have described the role of salt bridge formed between phosphorylated Ser188 of loop D and Lys270 of C-terminal of neighbouring monomer causing change in conformation of loop D which was earlier in closed form and leading to channel opening. In another study, it has been shown that phosphorylation of Ser111 of AQP4 doesn’t have any effect on conformation of Ser111 containing loop B or of the loop D as well as on water permeability (Assentoft et al., 2013). Rodrigues et al., used yeast system to show that AQP5 gating can be regulated by pH depending on phosphorylation (Rodrigues et al., 2016). They have suggested that at physiological pH condition i.e. 7.4, phosphorylated AQP5 induces the conformational changes at residue Ser and Thr facing cytoplasmic region. This leads to de-protonation of His183 residue facing outer membrane which leads to widening of the channel pore. Whereas at pH 5.1, the protonated residue and putative hydrogen bond interaction keeps the channel pore narrow but open resulting in lowering of the water permeability. There are numerous evidences of regulation of aquaporins, majority of these uses gating mechanism affecting the rate of water permeability dependent on phosphorylation. However, the possibility of AQP2 gating mechanism has not been rigorously explored.

AQP2 has been reported to be regulated by vasopressin-induced phosphorylation causing its trafficking to the apical membrane. AQP2 contains many putative kinase recognition residues such as Ser256, Ser261, Ser264, Ser269, and these C-terminal residues that are suggested to play important role in trafficking of AQP2 (Moeller et al., 2009a). Phosphorylation of Ser256 has been suggested to participate in AQP2 translocation to apical membrane but doesn’t affect water permeability through AQP2 channel (Brown et al., 2008). Phosphorylated AQP2 has comparable water permeability with that of wild type AQP2 whereas dephosphorylation of AQP2 reports threefold reduction in osmotic water permeability in comparison of wild type AQP2 (Moeller et al., 2009b). In another contrasting study by Eto et al, phosphorylated Ser256 has been shown to be involved in increased water permeability of AQP2 (Eto et al., 2010). Both groups have speculated that this phosphorylation may induce conformational changes in the C-terminal and are important for channel gating (Eto et al., 2010; Moeller et al., 2009b). This study was aimed at analysing the effect of phosphorylation of Ser256 of AQP2 on the conformation of AQP2 and its role in the regulation of water permeability. We investigated for the first time, the possible molecular basis of the effect of phosphorylation on water permeability of AQP2 and analyzed both the unphosphorylated and phosphorylated forms with the help of molecular dynamics simulations.

## Material and method

The unphosphorylated X-Ray diffracted crystal structure of’ human aquaporin-2 with resolution 3.05Å, has been obtained from the Protein Data Bank (PDB ID: 4OJ2) (Vahedi-Faridi et al., 2013). 3D structure was monomeric; thus, tetrameric form was constructed using online serverMakeMultimer.py (http://watcut.uwaterloo.ca/tools/makemultimer/index).

Crystal structure contains engineered amino acid at Ala 256 which was substituted with Ser256 identical to wildtype hAQP2. We were interested in studying effect of phosphorylation of Ser256 on water permeability. Therefore, phosphorylated form of AQP2 was generated by adding a phosphate group to Ser256 residue using CHARMM patch in NAMD (Phillips et al., 2005). Both unphosphorylated as well as phosphorylated systems were prepared by embedding in POPC lipid bilayer using MEMBRANE plugin of VMD (Humphrey et al., 1996). The system was then placed in TIP3P (Jorgensen et al., 1983) water box of size 130×130×100Å by using SOLVATE plugin of VMD. Water molecules were removed from hydrophobic regions of lipid bilayer while exposed part of AQP2 was surrounded by a layer of 20Å thick water molecules. Total number of water molecules in phosphorylated and unphosphorylated system was 28546 and 28502, respectively. Net charge of the system was made zero by adding appropriate number of counter ions using AUTOIONIZE plugin of VMD and ionic concentration was set to 150mM by adding Na^+^ and Cl^-^ions.

### MD simulations

Phosphorylated as well as the unphosphorylated structure of AQP2 were minimized in three stages Initially, both system were minimized for 8000 steps of steepest descent and then for 15000 steps of conjugate gradient. In third stage, system was minimized for 20000 conjugate gradient steps keeping the ar/R site residues fixed. Only lipid tails of minimized system were allowed to melt at 450K in NVT ensemble for 200 ps, while keeping rest of atoms (water, lipid head group, ions, and protein) fixed in order to introduce appropriate disorder of fluid like bilayer. The first equilibration was performed for 300ps at 310K by applying constraint of 4, 3, 2 and 5 kcal/mol/Å on lipid headgroups, tails, water, and protein, respectively. Later on, the system was equilibrated for 1000ps by reducing the constraint to 3, 2, 1 and 4 kcal/mol/Å. Again a constrained equilibration of 1000ps was performed applying force constant of 2, 1, 0, 2 and 4 kcal/mol/Å on headgroups, tails, water, protein and ar/R residues. The system was further equilibrated for 1.5ns with reduced constraint of 1kcal/mol/Å each on headgroups and tail while keeping the force constant of 2 kcal/mol/Å on protein and no constraint on the water molecules. An additional constraint of 4 kcal/mol/Å has been applied on dihedral angle of Arginine (Arg) residue of ar/R site to prevent initial protrusion and hence blocking of the pore. Further equilibration was carried out at 310K in NPT ensemble fixing only the protein atoms and dihedral angles of Arg with the constraint of 2 and 4 kcal/mol/Å, respectively for 3.5ns. Final equilibration of 7ns was performed with reduced constraint of 1.5 and 3 kcal/mol/Å on protein and dihedral angles of Arg, respectively in NPT ensemble. Later, all constraints were removed and the system was further simulated for 5ns in the same ensemble. Finally, simulation was carried out for both the phosphorylated and unphosphorylated systems for 100ns. NAMD software package (Phillips et al., 2005) was used for performing all MD simulations with CHARMM 27 parameter set (MacKerell et al., 1998). The pressure was set at 1atm using Langevin piston method (Feller et al., 1995) at a constant simulation temperature of 310K along with periodic boundary condition. Time step of 2fs and SHAKE algorithm (Ryckaert et al., 1977) were used and latter is being employed to apply a constraint on bonds involving hydrogen atoms. Particle Mesh Ewald (PME) method (Essmann et al., 1995) was used to calculate Electrostatic interactions. *Van der Waals* interactions were calculated interactions were calculated with switch distance of 10Å and cut off at 12Å .VMD was used for visualization and CPPTRAJ module (Roe and Cheatham, 2013) of Amber 16 was used to carry out MD trajectory analysis.

### Permeability coefficient

The single-file water molecules permeating through a channel was quantified using diffusion permeability coefficient (*p_d_*) given by

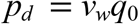

where *v_w_* is the volume of a single water molecule and *q*_0_ is the number of complete permeation events in one direction across the channel in unit time (Zhu et al., 2004). If the average number of water molecules in the lumen of the channel is *N*, the osmotic permeability constant (*p_f_*) can be calculated as

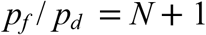

We have calculated the rate of diffusion permeability and osmotic permeability coefficient for each state of AQP2 i.e. phosphorylated and unphosphorylated through monomeric pore for 100ns of MD simulation run.

## Results

MD simulation trajectories of phosphorylated (phosphorylation at Ser256) and unphosphorylated AQP2 were analysed to observe any conformational changes in C-terminal region. The backbone root mean square deviations (RMSD) of transmembrane region of phosphorylated and unphosphaorylated AQP2 ranges between 1.48 to 2.09 Å and 1.42 to 2.16 Å, respectively. However, average RMSD of phosphorylated AQP2 (1.76 Å) is lower than that of unphosphorylated one (1.81 Å) depicting the stability of former structure (Fig 1). The variation of RMSDs of complete phosphorylated and unphosphorylated AQP2 is presented in S1 Fig. It has been observed that backbone RMSD of C-terminal region i.e. residues 225 to 257 ranges from 8 to 21Å and varies between 10 to 26Å, for phosphorylated and unphosphorylated AQP2, respectively (S2 Fig).

**Fig 1.**
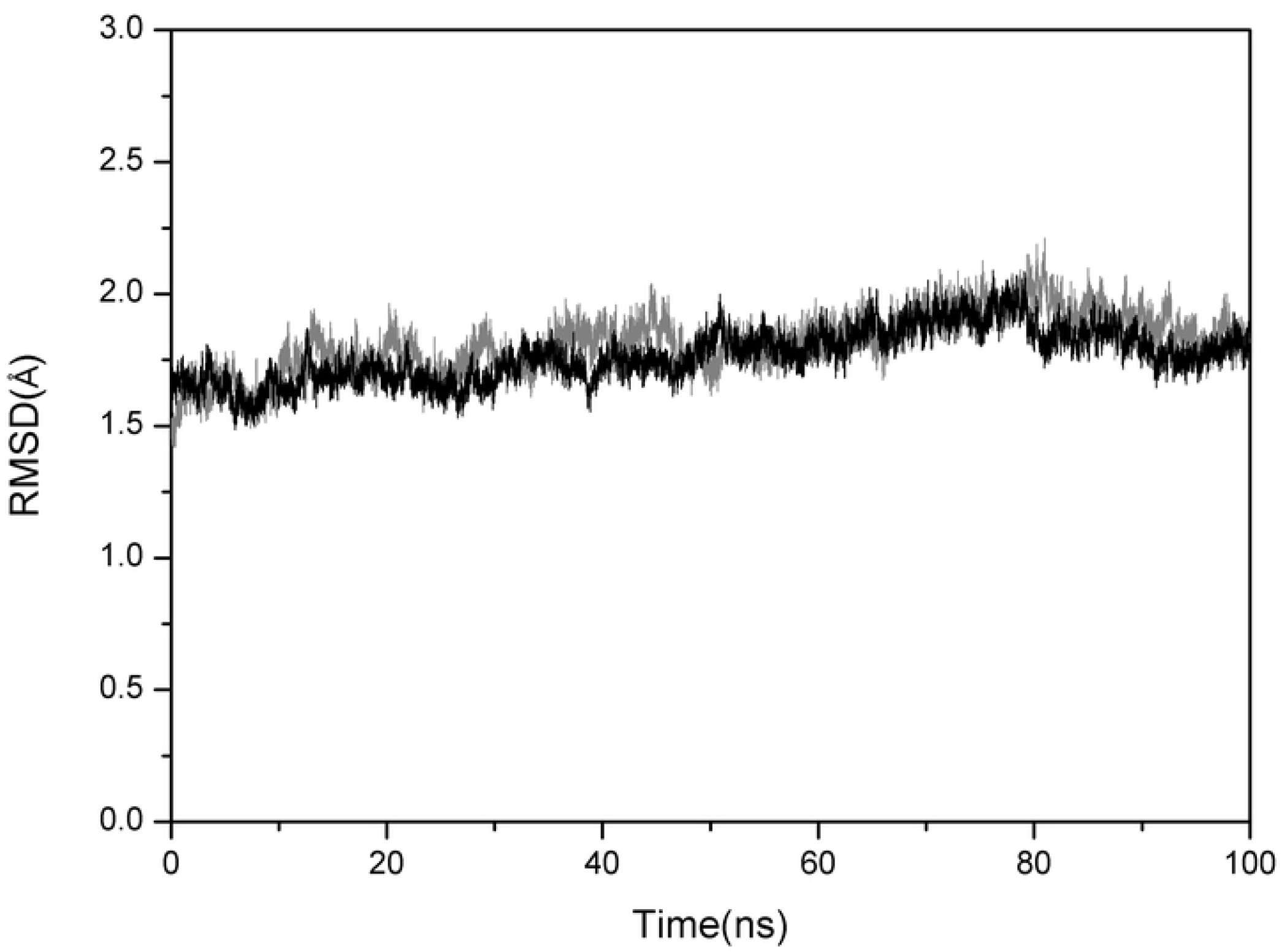
Backbone RMSD of the transmembrane region (1-224) of unphosphorylated (black) and phosphorylated (grey) AQP2 for 100ns of MD simulations

High RMSDs of C-terminal region possibly contribute to the high RMSD of the tetramer assembly of AQP2. Despite the overall high RMSD, the C-terminal region keeps its compactness which has been confirmed by calculations of Radius of Gyration (Rgyr) as shown in (Fig 2a, b, c and d). The values of Rgyr fluctuates between 13Å to 18Å for C-terminal region of phosphorylated as well as unphosphorylated monomer A and B of AQP2. On the other hand, Rgyr ranges between 11Å to 17Å for C-terminal region of monomer C and D during MD simulations in both forms of AQP2.

**Fig 2.**
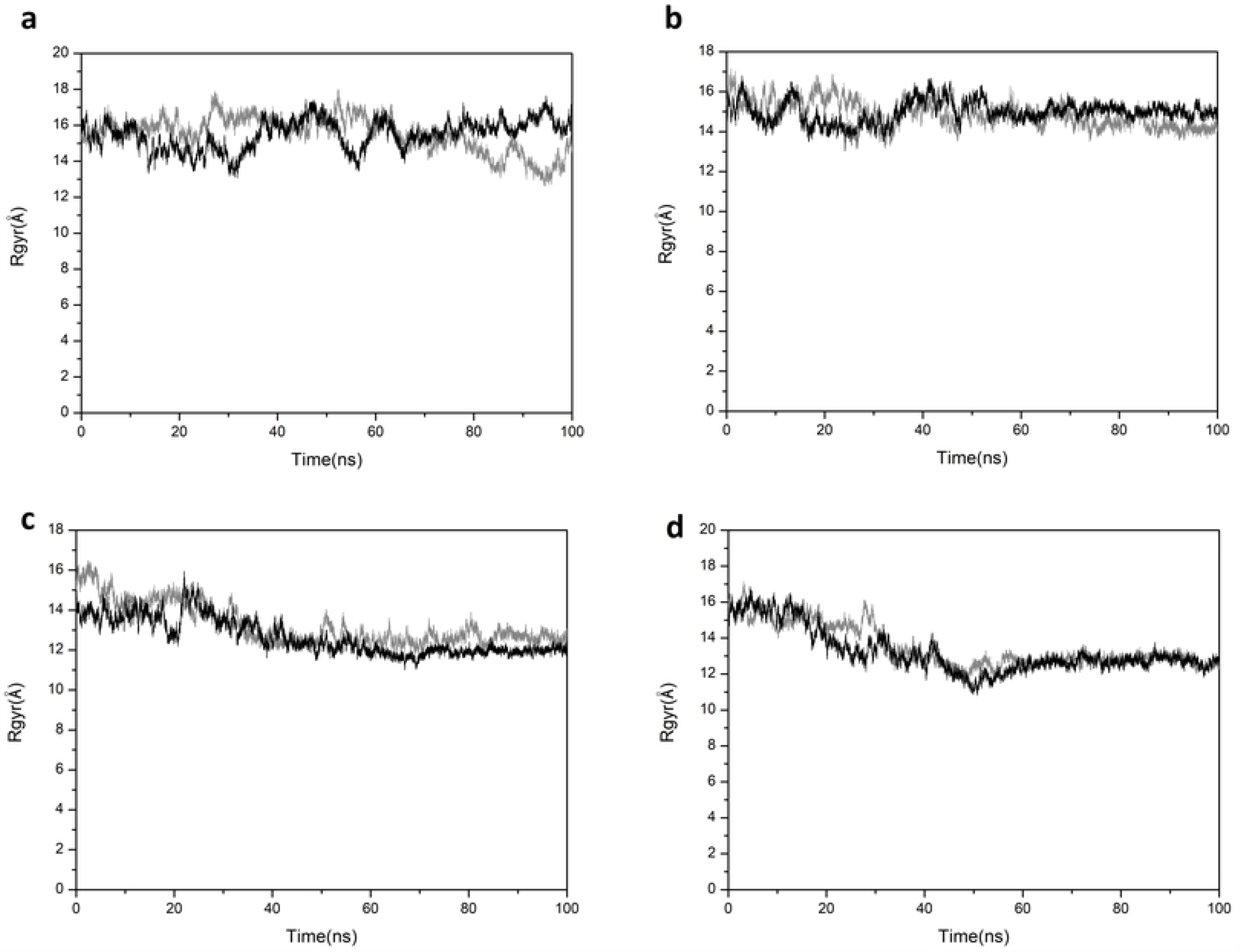
Radius of gyration of C-terminal region (225-257) of (a) monomer A (b) monomer B (c) monomer C (d) monomer D of AQP2 throughout 100ns MD simulations. Unphosphorylated and Phosphorylated AQP2 are represented by black and grey colours, respectively for each monomer of AQP2.

### Flexibility of C-terminal region of Phosphorylated vs Unphosphorylated APQ2

Root Mean Square Fluctuations (RMSFs) of individual monomers are high in C-terminal region (225 to 257) in comparison to the rest of the protein (Fig 3a, b, c and d). RMSF of amino acids of this C-terminal region increases up to 30 Å in case of monomer B of unphosphorylated AQP2 while for similar region of phosphorylated AQP2, it remains below 15Å.

**Fig 3.**
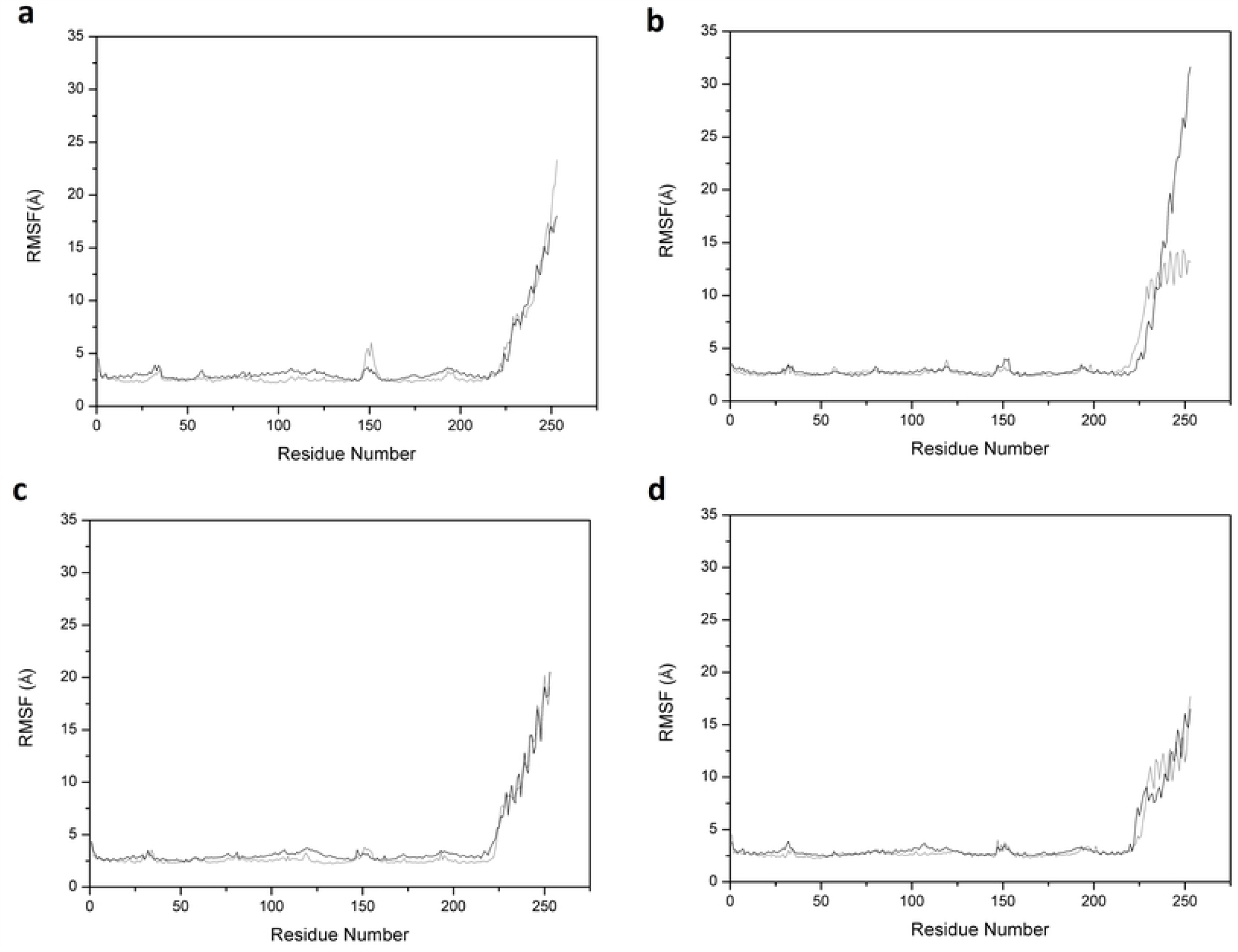
Root mean square fluctuations of (a) monomer A (b) monomer B (c) monomer C (d) monomer D of AQP2 over 100ns of MD simulations. Unphosphorylated and Phosphorylated AQP2 are represented by black and grey colours, respectively for each monomer of AQP2.

We have carried out disorder analysis of protein sequence using PrDOS (Ishida and Kinoshita, 2007) and disorder probability of each residue of AQP2 has been calculated. The prediction results showed that C-terminal region residues of AQP2 have high disorder propensity in comparison to rest of the protein (Fig 4a). High RMSFs in all the monomers of unphosphorylated as well as phosphorylated AQP2 are mainly attributed to the movement of C-terminal region present in the cytoplasmic side (Fig 4b).

**Fig 4.**
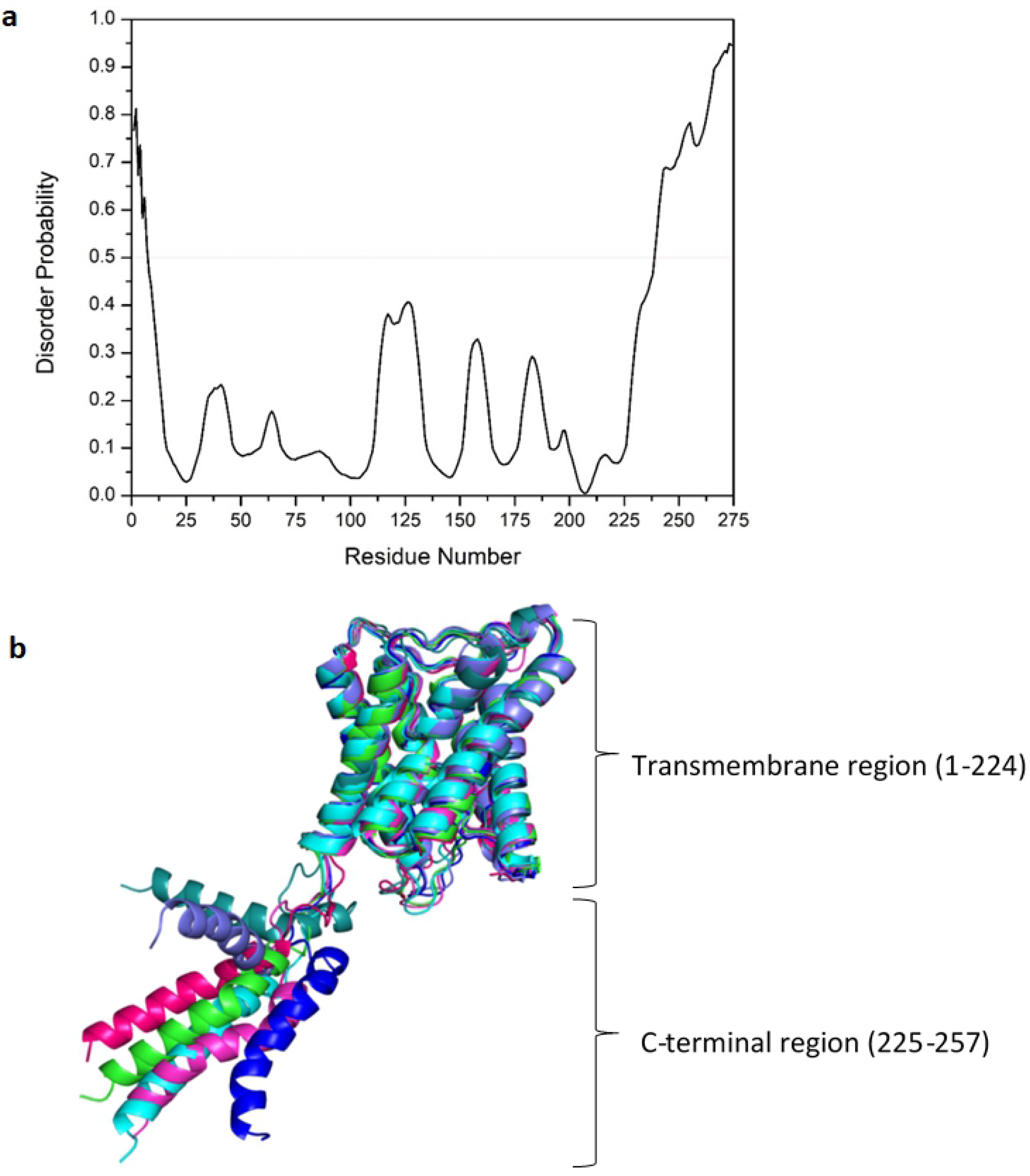
(a) Disorder Analysis of complete protein sequence using PrDOS; 0.5 is significance threshold for disordered residues (b) A few snapshots of monomer during MD simulations showing C-terminal flexibility and stable transmembrane. Various snapshots of monomeric AQP2 are shown in different colours across MD simulations.

A distinct fluctuation is observed from residue 150 to 155 in all monomers of AQP2, and this segment of protein comprises of a loop which is free to move around. These residues of monomers are also involved in intra and intermonomeric salt bridge formation. In unphosphorylated AQP2, intra-monomeric interactions between residues Glu155 & Arg153 of monomer C and Arg5 & Glu151 of monomer B were observed during MD simulations. Also, there are interactions between residues Asp150 & Arg152 of monomer C and those of monomer B, with the propensity of 99.65% and 93.43%, respectively. Inter-monomer interactions between Glu151 of monomer A and Arg153 of monomer C with occupancy of 56.51% has also been observed during the MD simulations. Whereas in case of phosphorylated AQP2, intra-monomer interaction between residue Glu155 and Arg153 of monomer B was found to have occupancy of 81.68% during the MD simulations. The occupancy of this interaction is higher than that in unphosphorylated AQP2 (54.1%). The interactions between Asp150 and Arg152 are found in both monomer A and monomer B of phosphorylated AQP2 with 80.23% and 100% occupancy, respectively. No inter monomeric interaction is found in this region of phosphorylated AQP2. The residues that are part of this region and show high fluctuations in RMSF are tabulated along with their interaction occupancies throughout MD simulations in Table 1.

**Table 1.**
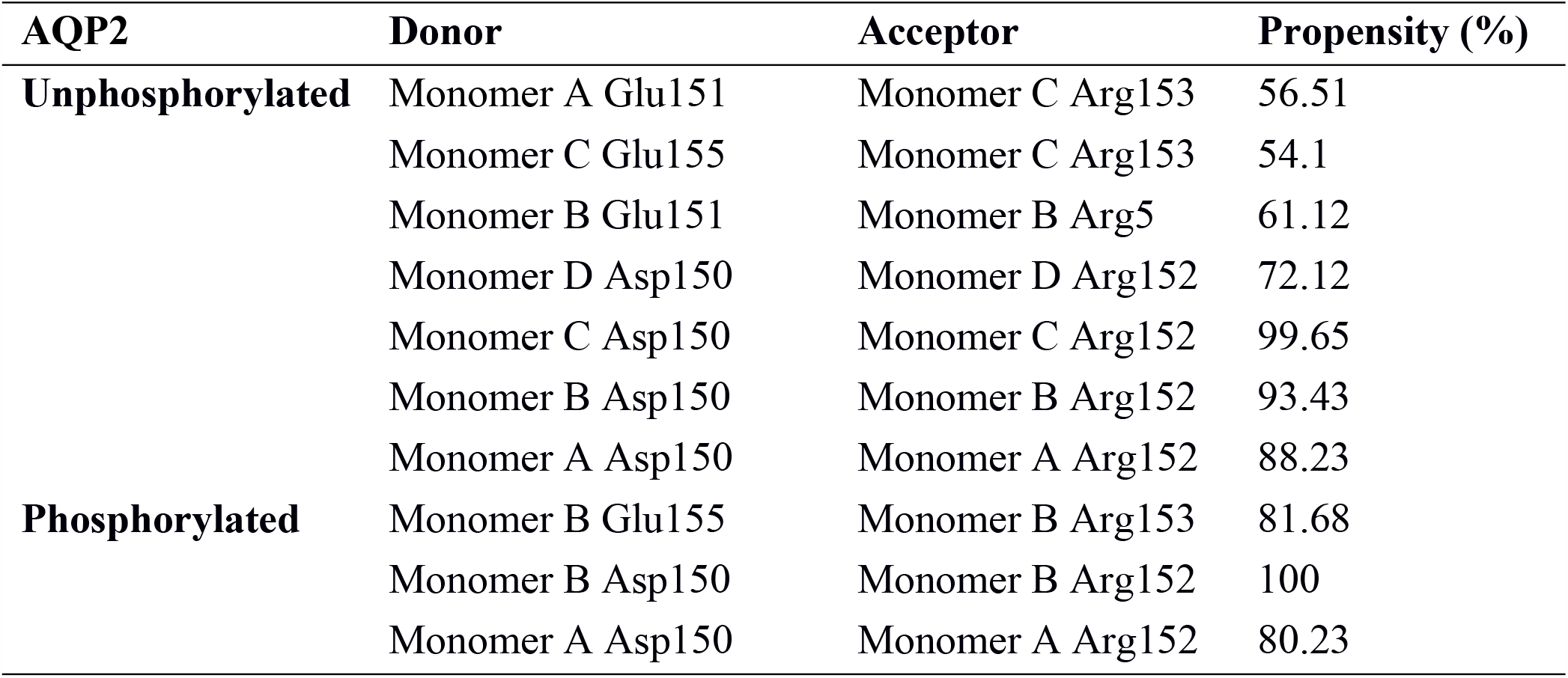
Occupancy of interactions between amino acid residues from 150 to 155 with higher RMS fluctuations in unphosphorylated and phosphorylated AQP2

### Intra monomeric interactions observed during MD simulations

Various intra monomeric interactions are found in AQP2 during MD simulations. Most of these are likely to be salt bridge interactions (Supplementary Table 1). Interactions with propensity of more than 35% are presented in this table. In unphosphorylated AQP2, there was an interaction between side chain oxygen of Glu250 and side chain nitrogen of Arg253 found in all three monomers except in monomer D with varying propensity of occurrence whereas, in phosphorylated AQP2, the interactions between these residues were absent in monomer C and have higher occupancy in monomer A, B, and D in comparison to unphosphorylated one. Interactions of amino acid residues with Asp245 in all the monomers have propensities lesser than 35% in case of phosphorylated in comparison to unphosphorylated AQP2. Propensity of interaction between Asp199 and Lys197 of monomer B of AQP2 has increased up to 70.9% in phosphorylated form (52% in unphosphorylated AQP2) and this interaction is reduced below 35% in case of monomer A of AQP2. Amino acid residues-Glu241 and Lys238 have shown interactions in all the monomers with different propensities in both phosphorylated and unphosphorylated forms. Most of the interactions shown in this table are consistent with those observed in the X-ray diffraction structure of the AQP2(Vahedi-Faridi et al., 2013). However, a few intra monomeric interactions having propensity above than 35% were observed only during MD simulations. In unphosphorylated AQP2, these interactions are between Glu248 and Lys228 of monomer C, between Glu232 and Arg153 of monomer C, between Asp245 and Arg252 of monomer A and between Asp 115 & Lys197 of monomer D. Interactions between Asp 199 and Lys197 were observed in all three monomers except in monomer D. In phosphorylated AQP2, interactions between Asp 115 and Lys 197 was found in monomer C and D with propensity of 48% and 49%, respectively.

### Interactions observed among neighboring monomers of AQP2

Diverse inter-monomeric interactions observed during MD simulations include interactions between the C-terminal regions of neighboring monomers and between the residues of N-terminal & C-terminal regions of monomers of AQP2. In the case of phosphorylated AQP2, there is an interaction between side chain oxygen of Ser256 of C-terminal of monomer D & side chain nitrogen of Arg5 of N-terminal of monomer B (Fig 5a). Also, there is interaction between oxygen of phosphate group of phosphorylated Ser256 of monomer D & side chain nitrogen of Arg5 of monomer B. These interactions appear to alter the conformation of highly flexible C-terminal of monomer D and displace this C-terminal of monomer away from the pore opening. The same phenomenon is not observed in the case of unphosphorylated one. Other interactions such as between side chain oxygen of Glu247 of monomer C & side chain nitrogen of Arg5 of monomer A (Fig 5b) and side chain nitrogen of Arg5 of monomer B with side chain oxygen of Glu248 of monomer D (Fig 5c) are found exclusively in phosphorylated AQP2. Interaction between side chain nitrogen of Arg5 of monomer D & side chain oxygen of Glu232 of monomer A is found in unphosphorylated form (Fig 5d). The latter interaction has not been observed in phosphorylated form of AQP2. A few inter-monomeric interactions with lower propensities were also observed and summarized in (Supplementay Table 2)

**Fig 5.**
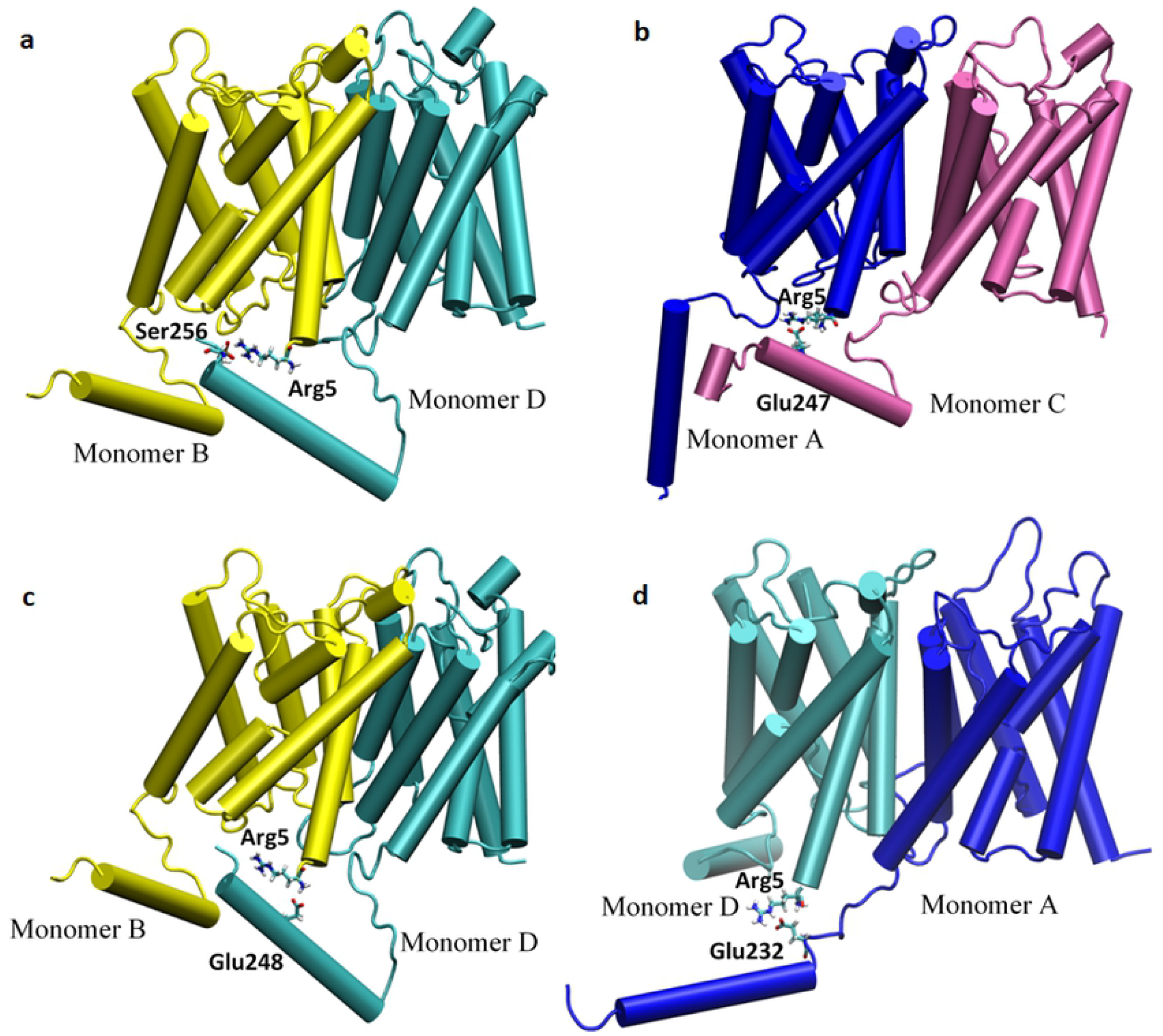
Interactions observed between neighbouring monomers of AQP2, in phosphorylated state (a, b, c), in unphosphorylated form (d)

### Conformational features of phosphorylated and unphosphorylated AQP2 monomeric pore

#### Electrostatic potential surface analysis

Snapshots of MD trajectories were extracted at an interval of 5ns from 100ns long MD trajectory. The electrostatic potential surface was analyzed using Pymol (Schrodinger, 2015). Surface representation of extracted structure at the start of MD simulations and at 25, 50, 75, 100ns for unphosphorylated as well as phosphorylated AQP2 are shown in Supplementary Fig 3. The regions surrounding the central and monomeric pore present positive electrostatic potential as shown by blue colour, whereas the region of C-terminal has majorly negative electrostatic potential shown by red color. Residues present surrounding the extracellular side of pore are mostly positively charged and hydrophobic amino acid. Residues present at cytoplasmic side or C-terminal region are mostly negatively charged and also aliphatic amino acid residues. There was no major difference in electrostatic potential of unphosphorylated and phosphorylated AQP2. The central pore was more clearly visible in phosphorylated AQP2 at various snapshots of the MD trajectory whereas monomeric pore were similar in both forms. Similar electrostatic potential depicts that the phosphorylation of C-terminal does not change the charge distribution of protein. Apart from showing electrostatic potential energy distribution, visualization of surface representation also corroborates that there was no marked occlusion of monomeric pore in either of the forms of AQP2. Structures of AQP2 at 50ns and 100ns of MD trajectory show that C-terminal is not able to cover the mouth of water channel in any of the forms (Fig 6 a, b, c, and d). Similar behaviour of conformations of AQP2 were found throughout the MD simulations.

**Fig 6.**
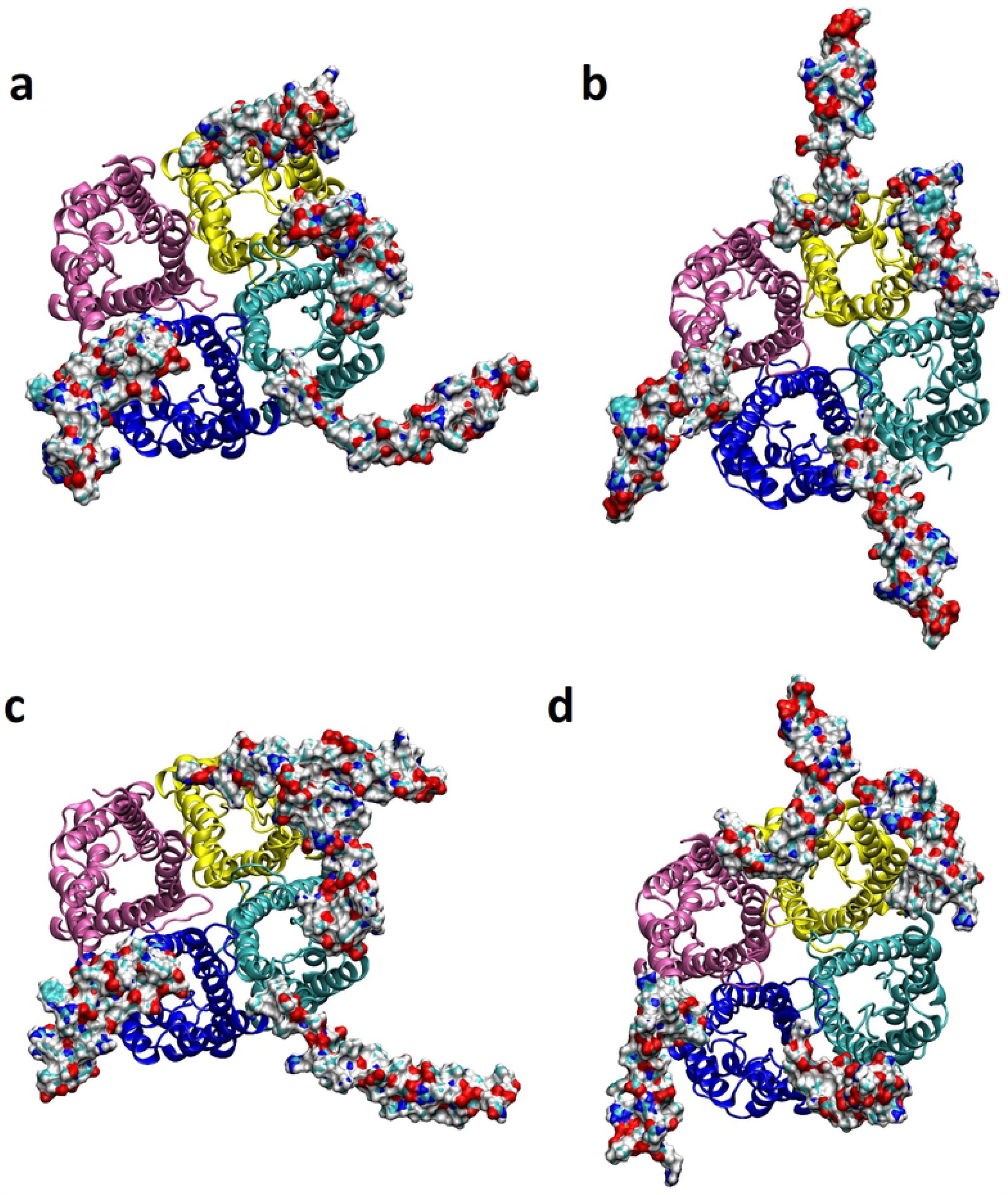
Surface representation of C-terminal region (225-257) showing snapshot at 50ns (a) unphosphorylated (b) phosphorylated AQP2 and at 100ns (c) unphosphorylated (d) phosphorylated AQP2 obtained from 100ns MD simulations. Amino acid residues for 1 to 224 of all monomers of AQP2 are shown in cartoon representation.

#### Hole Analysis

The pore radius of phosphorylated and unphosphorylated forms of AQP2 has been calculated during MD simulations using HOLE program (Smart et al., 1996). Analysis shows that there was a marginal difference in pore radius of two forms of AQP2 during MD trajectory. More precisely, the pore size may appear smaller in unphosphorylated form in comparison to phosphorylated AQP2, but there is no complete occlusion of any of the monomeric pores during 100ns of production run. The pore radius has been calculated at 5, 25, 50, 75, and 100ns of MD trajectory. There were discernable fluctuations in minimum pore radius values in all monomers of both forms of AQP2 and the change in pore radius along the channel axis has been shown in Fig 7. The pore radius calculation shows that the average minimum pore radius is as low as 0.23Å and as high as 4.98Å for unphosphorylated AQP2 and these values are 0.31Å and 4.95Å as lowest and highest, respectively in phosphorylated AQP2. The minimum size of pore radius of phosphorylated AQP2 is slightly higher throughout the MD trajectory in comparison to unphosphorylated form and allows free movement of water molecules through the channel. In both phosphorylated and unphosphorylated AQP2, the respective lowest pore radii were observed at ar/R constriction site and lumen region comprising NPA motifs. The latter region acts as a selectivity filter and plays a crucial role in water transport across the channel. Pore radius analysis of duplicate MD simulations for 100ns, also shows marginal difference in pore radius of unphosphorylated AQP2 in comparison to phosphorylated AQP2.

**Fig 7.**
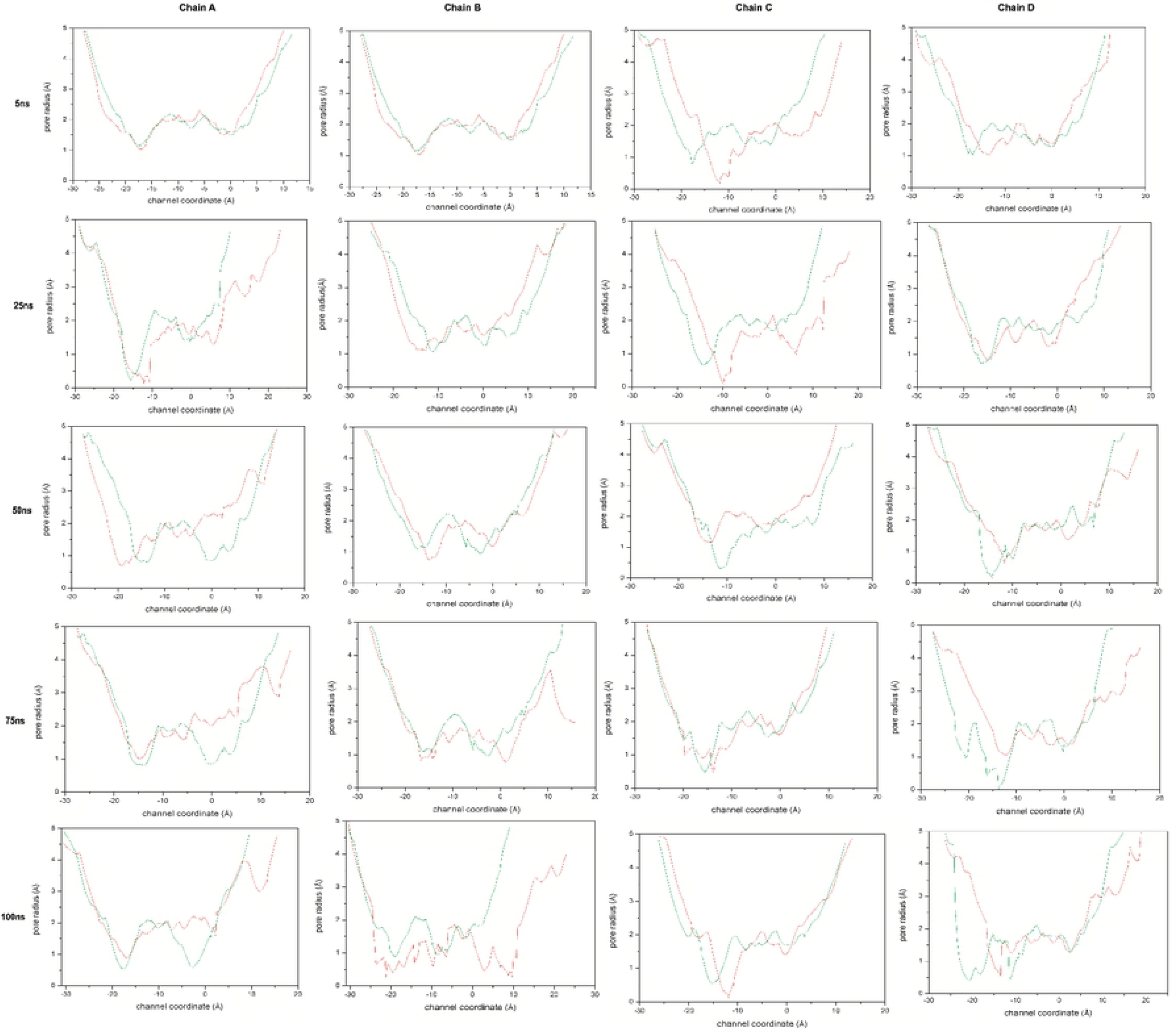
Hole analysis of pore radii of unphosphorylated AQP2 (red) phosphorylated AQP2 (green) showing changes in pore radius across the channel coordinate at 5ns,25ns,50ns,75ns, and 100ns MD simulations.

#### Effect of phosphorylation on water conductivity

The calculated diffusion permeability of unphosphorylated and phosphorylated AQP2 was 0.63×10^−14^ cm^3^ s^−1^ and 0.74×10^−14^ cm^3^ s^−1^ respectively, while osmotic permeability was 4.96×10^−14^ cm^3^ s^−1^ and 5.92×10^−14^ cm^3^ s^−1^ for unphosphorylated and phosphorylated AQP2, respectively. These values show little difference in the rate of diffusion and osmotic permeability in two forms of AQP2. The trajectories of both the systems were visualized using VMD showing the movement of water molecules through the channel during MD simulation run. There was a continuous flow of water across the channel irrespective of phosphorylated or unphosphorylated forms of AQP2. We have shown single file water flow through monomer A and B of unphosphorylated and phosphorylated AQP2, similar condition is observed in other monomers (Fig 8a and b).

**Fig 8.**
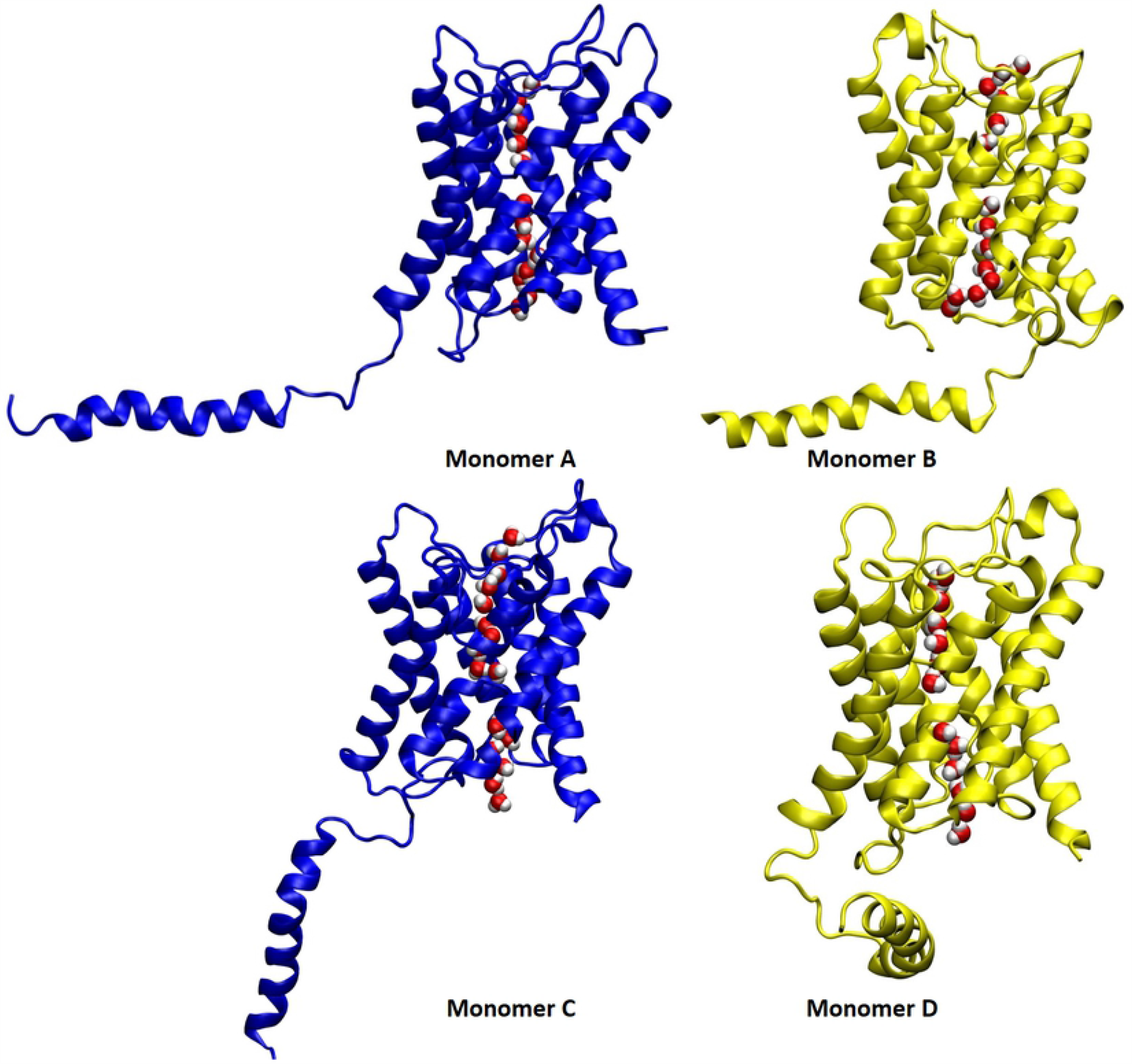
Snapshot of the last structure of MD trajectory showing a continuous flow of water across the monomeric channel in (a) unphosphorylated and (b) phosphorylated AQP2. Water is shown in VDW and AQP2 monomers are shown in cartoon representation.

## Discussion

Till date, various experimental studies have been carried out to observe the effect of phosphorylation on the AQP2 transport to apical membrane and on water permeability (Eto et al., 2010; Noda and Sasaki, 2006; Xie et al., 2010). Osmotic gradient is the driving factor for water transport through aquaporin water channel across the epithelial plasma membrane (Walz et al., 1997). Water enters the principal cell of collecting duct via apical AQP2. The majority of AQP2 remains localized in the subapical vesicle in the absence of arginine vasopressin (AVP), and results in low water permeability of principal cells (Nielsen et al., 1993). AVP binds to the AVP receptor initializing a signal cascade wherein the level of cAMP increases, subsequently Protein kinase A (PKA) gets activated. PKA activation leads to phosphorylation of AQP2 which is probably an important phenomenon of AQP2 trafficking (Kuwahara et al., 1995). There are four Serines (Ser) Ser256, Ser261, Ser264 and Ser269 located in the cytoplasmic C-terminal region of AQP2 and these undergoes phosphorylation or dephosphorylation in response to vasopressin. Among these serines, Ser256 is considered to be critically important residue that undergoes phosphorylation and triggers the accumulation of AQP2 in the plasma membrane (Fushimi et al., 1997). Apart from trafficking mechanism, there have been studies that show the relationship between phosphorylation and gating mechanism in aquaporins. There are various examples corroborating aquaporin gating, especially in plant aquaporins. The spinach aquaporin SoPIP2;1 is one of the best illustrations of phosphorylation induced gating. MD simulation shows that there was interaction between Ser188 and Lys270 of the C-terminal of SoPIP2;1 causing conformational changes in loop D and leading to opening of channel (Nyblom et al., 2009). A study by Hiroaki et al. suggested that phosphorylated Ser180 may bind to C-terminal domain and could block the cytoplasmic entrance of AQP4 water channel (Hiroaki et al., 2006). However, this was not supported by a molecular dynamics study by Sachdeva and Singh showing that there was marginal effect of phosphorylation of Ser180 on water permeability of AQP4 (Sachdeva and Singh, 2014). It has also been established experimentally by Assentoft et al. that AQP4 plasma membrane trafficking and channel gating were not significantly regulated by phosphorylation of serine at COOH terminal (Assentoft et al., 2014). There have been a few studies supporting the theory that Ser256 phosphorylation has direct involvement in increasing the water permeability of AQP2 and could bring some conformational changes in C-terminal important for channel gating as speculated (Eto et al., 2010; Fushimi et al., 1997; Moeller et al., 2010). Kuwahara et al. showed that phosphorylation of individual AQP2 increases the water permeability of AQP2 expressed in Xenopus oocytes although the amount of AQP2 on oocytes didn’t increase (Kuwahara et al., 1995). In contrast to this, Fushimi et al. showed that there was a good correlation between number of AQP2 water channels and increase in permeability of wild type as well as mutant (S256A) AQP2 rather than phosphorylation of individual AQP2 channel (Fushimi et al., 1997). On the contrary, Lande et al. used purified endosome from the apical membrane of rat inner medullary collecting duct cells that are highly enriched for AQP2 and showed that water permeability is not affected by AQP2 phosphorylation (Lande et al., 1996). Kamsteeg et al. and Moeller et al. also demonstrated oocytes expressing S256D and wild type AQP2 in equal abundance on plasma membrane have similar water permeability and they also excluded the involvement of phosphorylation dependent channel gating on regulation water permeability of AQP2 (Kamsteeg et al., 2000; Moeller et al., 2009b). But due to unavailability of AQP2 crystal structure (till 2014), these studies could not exclusively validate or invalidate the theory that C-terminus phosphorylation mediated AQP2 channel gating has role in water permeability regulation in AQP2. Hence, structural analysis was much needed to confirm the possibility of channel gating because of possible conformational change in C-terminal residue (exclusively Ser256) due to phosphorylation and also its effect on water permeability of AQP2. In present molecular dynamics study, we have shown that phosphorylated AQP2 has overall more stable structure throughout MD simulations in comparison to unphosphorylated. RMSD values for unphosphorylated and phosphorylated AQP2 are high because of highly mobile C-terminal region whereas transmembrane region does not show much mobility and has low RMSD values regardless of phosphorylation of AQP2. In tetramer assembly of AQP2, each monomer had unique intra-as well as inter-monomeric interactions and function independently as individual water channel (Gomes et al., 2009). Therefore, it becomes necessary to study the tetrameric form of AQP2. The inter-monomeric interaction between Ser256 of C-terminal of monomer D and arginine of N-terminal of monomer B of phosphorylated AQP2 seems to be involved in displacing the C-terminal residues outward or away from pore formed by monomer D so that it could not occlude the water channel. This interaction was not observed in unphosphorylated form and the C-terminal was free to move towards the pore of the water channel at a few times in MD trajectory. In phosphorylated AQP2, all the monomeric chains were involved in inter-monomeric interactions but this is not same in case of unphosphorylated AQP2 which shows interaction between monomer D and A. Surface electrostatic potential evaluation by Oliva et al. showed negative values for cytoplasmic half and positive for extracellular half of AQPs, a similar trend in orthodox AQPs (Oliva et al., 2010). In our study, similar electrostatic potential profile was observed in unphosphorylated as well as phosphorylated AQP2 during MD trajectory. The surface representation at the different time points of trajectory for both the forms show that there is no pore occlusion at the starting structure and the C-terminal regions are away from the monomeric pore of water channel but as the MD simulation proceeds, expected movement in C-terminal was observed. In phosphorylated AQP2, there was a slight movement of C-terminal region of monomer B towards its own center of pore channel and but it doesn’t occlude the pore. We have also observed that C-terminal of monomer C also moves slightly near to the mouth of channel formed by monomer B and C-terminal of monomer D moves towards monomeric pore formed by monomer A. These movements do not occlude the monomeric pore of the channel of phosphorylated AQP2. There is a continuous flow of water through the channels of these monomers. In unphosphorylated AQP2, C-terminal region of monomer B also moves near the pore of monomer D while C-terminal of monomer D moves towards mouth of monomeric channel formed by monomer B. C-terminal of monomer C traverses to the mouth of water channel of monomer A of AQP2. It appears to marginally reduce the pore radius in the case of unphosphorylated AQP2, however, it doesn’t obstruct the pore completely and hence the continuous flow of water molecule was not hindered. The change in pore radius has also been followed up using hole analysis which depicts that the radius of pore remains same in phosphorylated as in unphosphorylated AQP2 in the beginning of MD simulations. Towards the end of MD simulations, there was slight reduction in pore radii specially in monomer B, C, D of unphosphorylated AQP2. However, this decrease was not capable of impeding the water flow through channel. The difference in water permeability calculated for phosphorylated and unphosphorylated AQP2 is negligible. These findings suggest that phosphorylation of Ser256 and resulting interactions as well as conformational changes in C-terminal may not be directly involved in regulation of water permeability and channel gating.

## Conclusion

Phosphorylation of Ser256 of C-terminal of AQP2 has been suspected to play role in its either trafficking or gating of the water channel. We have carried out molecular dynamics simulation study to explore the possibility of involvement of Ser256 phosphorylation in gating of the channel. The study shows that AQP2 has highly flexible C-terminal region (225-257) but it is slightly stable in case of phosphorylated AQP2, possibly due to interactions occurring particularly in phosphorylated and not in unphosphorylated AQP2. The interactions between Ser256 of C-terminal of monomer D & Arg5 of N-terminal of monomer B, between Glu247 of monomer C & Arg5 of monomer A and Arg5 of monomer B with Glu248 of monomer D were found exclusively in phosphorylated AQP2. Hole analysis of pore radius of AQP2 shows marginal difference in pore sizes of unphosphorylated and phosphorylated AQP2. The water permeability calculation for unphosphorylated AQP2 (4.96×10^−14^ cm^3^ s^−1^) was marginally low than the phosphorylated AQP2 (5.92×10^−14^ cm^3^ s^−1^). The conformational analysis of both the forms of AQP2 also shows that there was no significant occlusion of pore upon phosphorylation of Ser256 and no other channel blocking residues were observed during MD simulations. Thus, these findings suggest that phosphorylation of Ser256 of C-terminal region may not contribute in the gating of AQP2. On the other hand, involvement of Ser256 phosphorylation in trafficking of AQP2 can not be ruled out as proposed in other studies.

## Supporting information

**S1 Fig**. Backbone RMSD of phosphorylated (grey) and unphosphorylated (black) AQP2 for 100ns of production run.

**S2 Fig**. Backbone RMSD of only C-terminal (225-257) region of phosphorylated (grey) and unphosphorylated (black) AQP2.

**S3 Fig**. Surface representation showing electrostatic potentials during 100ns of MD simulations of AQP2. Electrostatic surface potentials were colored red and blue for negative and positive charges, respectively and white color represents neutral residues. In this Fig, (a) shows the surface electrostatic potential of starting structure of unphosphorylated and phosphorylated AQP2. Likewise, (b)-(e) shows structures at 25ns, 50ns, 75ns and 100ns respectively. Left panel shows unphosphorylated and right one shows phosphorylated AQP2

**S1 Table**. Propensities of intra monomeric interactions (salt bridge) in unphosphorylated and phosphorylated AQP2.

**S2 Table**. Propensities of inter monomeric interactions (salt bridge) in unphosphorylated and phosphorylated AQP2.

## Acknowledgment

Ms Pragya Priyadarshini acknowledges the award of DST-INSPIRE fellowship from Department of Science and Technology, New Delhi. Authors are thankful for support from Council of Scientific and Industrial Research and Department of Biotechnology, New Delhi, India.

